# Frequency-dependent Inhibition during Deep Brain Stimulation of Thalamic Ventral Intermediate Nuclei

**DOI:** 10.1101/2025.09.10.675201

**Authors:** Zoe Paraskevopoulos, Yupeng Tian, David Crompton, Suneil K. Kalia, Mojgan Hodaie, Andres M. Lozano, Luka Milosevic, William D. Hutchison, Milad Lankarany

**Author notes:** Correspondence to: Milad Lankarany Krembil Brain Institute, University Health Network 60 Leonard Ave, 4KD512, Toronto, ON M5T 0S8, Canada. These authors contributed equally to this work (Zoe Paraskevopoulos, Yupeng Tian).

## Abstract

Deep brain stimulation (DBS) of the thalamic ventral intermediate nucleus (Vim) has been a standard therapy for essential tremor. It has been shown that high frequency (≥100Hz) DBS suppresses Vim neuronal firing and tremor activity, however, the underlying mechanisms are not fully understood. Here, we use in vivo recordings (single-unit) of Vim neurons (n=19, people with essential tremor) during different DBS frequencies to investigate whether neuronal suppression during high-frequency DBS occurs at synaptic/cellular levels (e.g., cell inhibition due to synaptic depression/fatigue during high-frequency DBS) or is influenced by network-level effects (e.g., recurrent inhibition). We propose a theoretical framework that explains DBS effects at both cellular and network levels, i.e., (continuous) high-frequency DBS not only depresses synapses projecting to Vim but also enables the recruitment of inhibitory neurons. A transient burst in the spiking activity of Vim during high-frequency DBS, prior to neuronal suppression, is likely providing sufficient network engagement to recruit inhibitory neurons that are silent during low-frequency DBS. Further, we detected a positive-going evoked-field potential effect, hereafter referred to as *quasi-evoked inhibition*, during high-frequency (100 Hz and 200 Hz) Vim-DBS in four out of 19 recording sites.

Interestingly, it was observed that (***i***) neuronal suppression is stronger in these four neurons (*P* < 0.05), implying that inhibitory engagement during high-frequency DBS can further suppress neuronal firing, and (***ii***) quasi-evoked inhibition emerges after the transient burst (*P* < 1.00 x 10^-7^), i.e., the latter may give rise to the former. By removing DBS artifacts with a novel algorithm and characterizing the dynamics of quasi-evoked inhibitory activity, we showed that the likelihood of occurrence of this inhibitory activity negatively correlated with the instantaneous firing rate (*P* < 1.00 x 10^-5^). These results suggest that an excitatory-inhibitory balance is likely regulating Vim activities during high-frequency DBS. Our findings shed light on potential network mechanisms underlying Vim-DBS, which can provide insight for optimizing DBS by designing new stimulation patterns.

## Introduction

Essential tremor (ET), regarded as a disorder of the cerebellum,^1–3^ is the most common tremor disorder affecting 5.79% of adults over 65 worldwide.^4^ It involves progressive kinetic tremor of the arms.^5,6^ and sometimes additional tremor affecting the head,^7^ gait abnormalities,^8^ ataxia,^9^ cognitive impairments,^10^ and personality disturbances,^11^ leading to overall decline in quality of life.^12^ While some studies found some neurodegenerative changes like Purkinje cell loss in ET,^13–17^ neurophysiological studies showed a reduction of GABA-A and GABA-B receptors compared to control subjects.^18^ The lack or reduction of inhibition (e.g., in Purkinje cells) may lead to propagation of overactivity (disinhibition) through the cerebello-thalamo-cortical network. Vim neurons in the thalamus, main targets of invasive deep brain stimulation (DBS),^19–23^ are thought of as relays that pass oscillatory tremor-related activity from the cerebellum to the cortex.^24,25^ High frequency (≥100Hz) DBS suppresses Vim’s neuronal firing and tremor severity.^26–28^ Despite clinical efficacy, the mechanisms of HF-DBS in Vim, whether neuronal suppression is a result of synaptic-cellular depression or network inhibitory inputs, are not fully uncovered.

One hypothesis for the mechanism of HF-DBS is that it leads to the recurrent activation of targeted synapses resulting in vesicle depletion.^29–30^ Vesicle depletion results in synaptic fatigue, and an overall reduced post-synaptic firing rate is observed because vesicle resources are limited for replenishment.^29–31^ Both experimental and computational research have shown support for this hypothesis.^28,29,32^ Erez et al. (2009) observed during HFS of the primate internal globus pallidus that a time-locked response occurred along with reduced firing rate.^32^ This response ceased after stimulation, implying short-term depression.^32^

Another hypothesis to explain the mechanism of HF-DBS is that it allows for the recruitment of inhibitory responses, leading to suppression of the target.^33^ Steiner et al. (2024) conducted human single-unit extracellular potential analysis of positive-going evoked potentials, suggesting that during HF-DBS of the subthalamic nucleus (STN), the external globus pallidus (GPe) is continuously activated both orthodromically and antidromically by the STN, resulting in increased net inhibitory feedback to the STN.^34^

In this work, we investigated both mechanistic theories of DBS in human Vim, leveraging single-unit extracellular spiking and field potential recordings from previous work.^28,35,36^ Using sophisticated computational modeling and model fitting,^37^ we have recently shown that during HF-DBS, the strength of inhibitory recurrent feedback to the Vim must be substantially higher than that during other (lower) frequencies to replicate experimental results reliably.^37^

The Vim is an excitatory thalamic nucleus^28^ that projects to cortical motor regions^28,38–40^ and thalamic interneurons.^41^ It also receives direct excitatory glutamatergic afferents from the cerebral cortex^11,42–44^ and the dentate nucleus of the cerebellum (Fig. 1a).^41,45–47^ The direct inhibitory inputs to the Vim come from GABAergic thalamic reticular projections^11,47–49^ and thalamic interneurons (Fig. 1a).^41,50^ If suppression of the Vim during HF-DBS occurs due to network inhibition, it may be due to orthodromic activation of thalamic interneurons, resulting in an increased inhibitory recurrent response. Another possibility is that HF-DBS of the Vim may activate the cortex both orthodromically and antidromically, resulting in increased activation of inhibitory thalamic reticular projection (TRN) efferents to the Vim.

**Figure 1.**
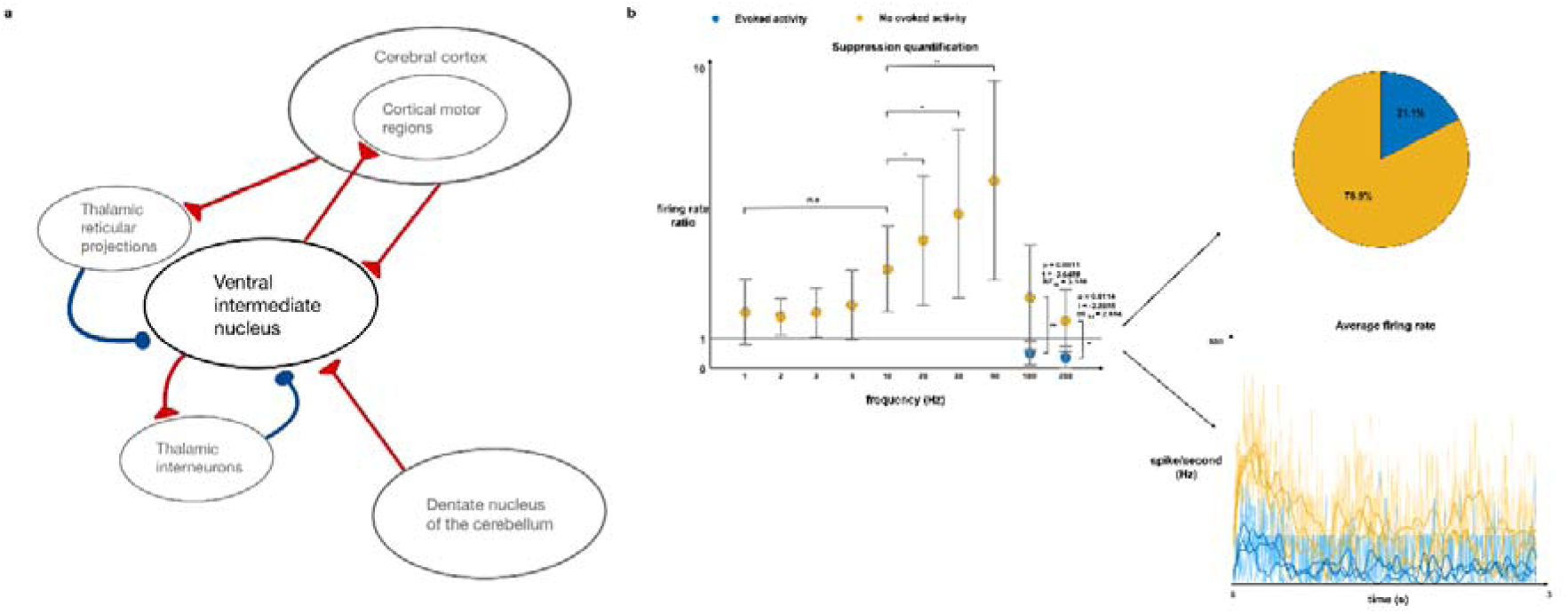
Stronger suppression following HFS in neurons with quasi-evoked inhibition and underlying circuitry: Subfigure **a** shows the direct afferent signals that project onto the Vim as well as efferent outputs. Excitatory signals are depicted in red and inhibitory signals in blue. The left of subfigure **b** plots the group average ratio between baseline firing rate and steady state firing rate at different DBS frequencies. At each point the standard deviation is also plotted. The yellow points include the group average ratio for neurons with no quasi-evoked inhibition and the blue points include the group average ratio for neurons with quasi-evoked inhibition. A horizontal line (black) is plotted at y = 1 to separate an increase in firing from baseline (above) from a reduction in firing from baseline (below). Test statistic and P-value results are shown for 100 Hz and 200 Hz. On the right of subfigure **b**, a pie chart shows the percentage of neurons evaluated with (blue) and without (yellow) quasi-evoked inhibition, and beneath a plot showing the average firing rate of three samples with quasi-evoked inhibition (blue) and three samples without quasi-evoked inhibition (yellow) are shown. The darker lines are the average firing rates using a 25 ms kernel and the lighter lines are using a 2.5 ms kernel. Left-tailed independent samples t-test: *\*P-value < 0.05; ** P-value < 0.005.* Wilcoxon rank sum test: *\*\* P-value < 0.005*.

Based on evidence supporting both potential mechanisms of DBS, we hypothesize that during HF-DBS of the Vim, the initial excitatory burst results in both synaptic depression and allows for inhibition recruitment, leading to overall suppression at the cellular and network levels. Additionally, we hypothesize that quasi-evoked inhibition, a positive-going evoked potential observed in a subset of Vim single-unit recordings (Fig. 2), is a marker of inhibition from the network. To explore these ideas, we developed a novel DBS artifact removal to preserve the shape of the quasi-evoked inhibition event; detected and temporally evaluated the amplitude of the quasi-evoked inhibition event; correlated quasi-evoked inhibition with firing suppression both within recordings, and between recordings using neurons that do not display this effect as a control group.

**Figure 2.**
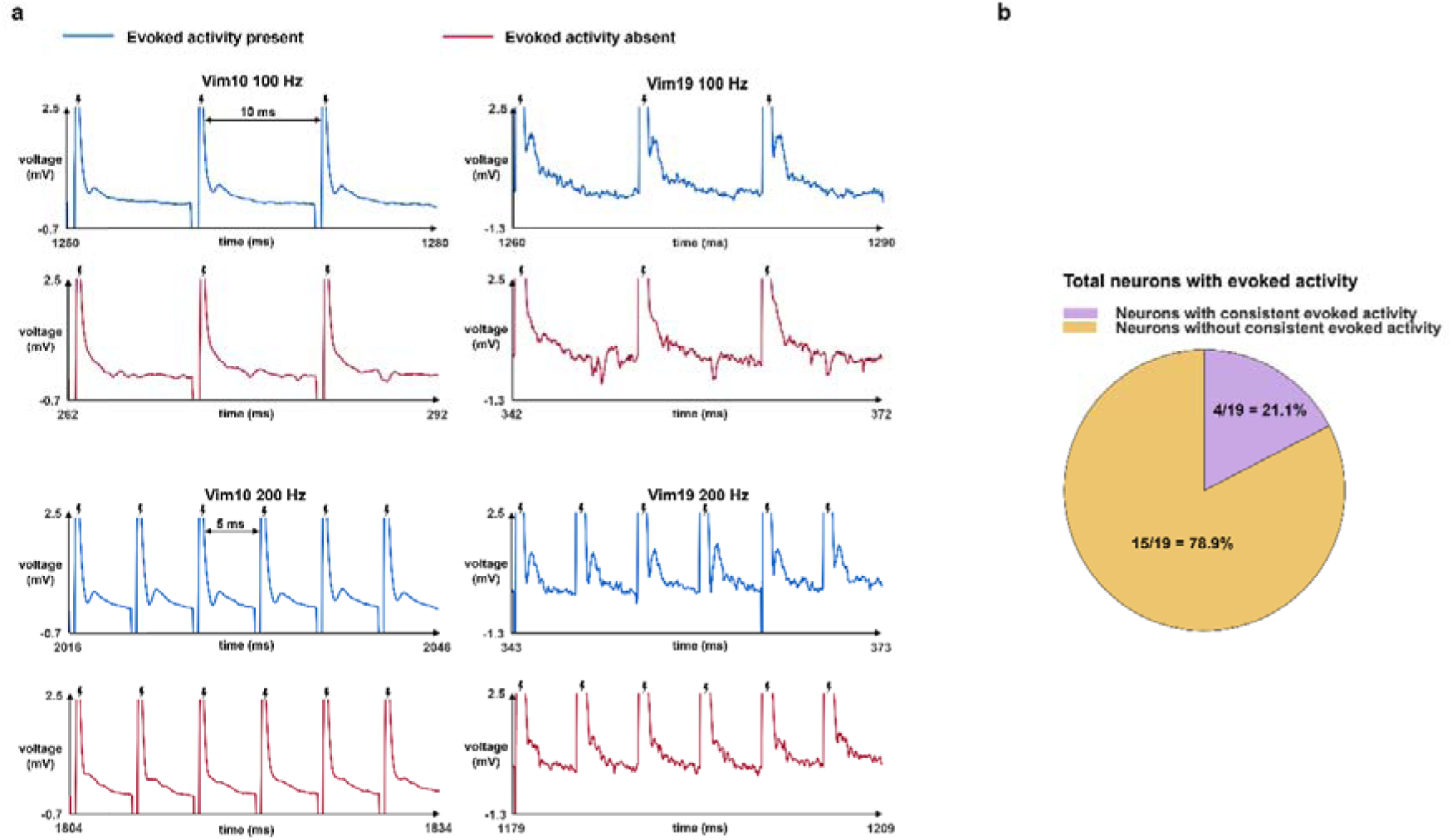
Quasi-evoked inhibition is observed in a subset of Vim neurons following DBS: In vivo human extracellular electrophysiological recordings are measured over time. 30 ms samples are shown from 3 s recordings in two different Vim neurons at both 100 and 200 Hz DBS frequency. The blue recordings represent areas of the recording with quasi-evoked inhibition present following the DBS pulse (marked with black icon). The pink recordings represent another area of the same recording where no quasi-evoked inhibition following the DBS pulse can be observed. The pie chart demonstrates the percentage of 19 total neurons, at greater or equal to 100 Hz DBS frequency, that consistently show quasi-evoked inhibition. Four neurons out of 19 or 21.1% demonstrate consistent quasi-evoked inhibition at HFS and 15 out of 19 or 78.9% do not.

## Materials and methods

### Human experimental data

We used the same human experimental single-unit recordings which were previously published in Milosevic et al. (2021).^35^ The commitment to ethics policies were previously fulfilled. All human experiments conformed to the guidelines set by the Tri-Council Policy on Ethical Conduct for Research Involving Humans and were approved by the University Health Network Research Ethics Board.^35^ Additionally, before the study, each patient provided written informed consent.^35^ The human experimental data protocols were from Milosevic et al. (2021).^35^ Microelectrodes were used to both deliver DBS and perform single-unit recordings. DBS was delivered using 100 µA and symmetric 0.3ms biphasic pulses (150µs cathodal followed by 150 µs anodal), recordings were sampled at >=10 kHz.

In this work, we used the outlined previously acquired^28,35^ and freely accessible^36^ data of single-unit extracellular recordings of spiking activity and field potentials in the thalamic ventral intermediate nucleus (Vim) of people with essential tremor, during various DBS frequencies (1-200 Hz) in Vim. The single-unit recordings were between 1-10 s long for frequencies 1-50 Hz and either 3 or 5 s long for 100 and 200 Hz; for each frequency of DBS, we analyzed eight to 18 recordings in different patients (total number of patients = 19). To localize spike timings from the single-unit recordings, we applied spike template matching. For each single-unit recording, a median filter was applied to remove baseline fluctuations and then spikes were identified by template matching using Spike2 (Cambridge Electronic Design, UK). Once this was completed, each file was manually inspected in 40 ms windows across the entire recording to verify the accuracy of spike detection.

### Artifact removal

The stimulation artifact is a result of resistance-capacitance effects in the system.^51–53^ The quasi-evoked inhibition events were merged with the DBS pulse artifacts, which need to be removed before analyzing the evoked temporal characteristics. Additionally, we wanted to exclude quasi-evoked inhibition events with very low intensities, as this was observed in Vim02 at 30-50 Hz stimulation. To threshold the quasi-evoked inhibition events and evaluate their characteristics, we developed a novel artifact removal to apply to the raw electrophysiological recordings. To preserve the features of the quasi-evoked inhibition events, we performed the artifact removal process in three steps (Fig. 3a, Supplementary methods). In the steps that did not involve preservation of the quasi-evoked inhibition, we used exponentials as they represent the summed decay of the resistance-capacitance properties.^51–53^

1. Apply a three parameter single exponential fit from the peak of the DBS pulse artifact to the start of the quasi-evoked inhibition event. This was subtracted from the original signal.
2. Apply a linear fit from the start to the end of the quasi-evoked inhibition event. This was done to prevent the quasi-evoked inhibition from being corrupted by the curvature of a nonlinear fit. It was subtracted from the original signal.
3. Apply a three parameter single exponential fit from the end of the quasi-evoked inhibition event to the end of the interpulse interval (IPI). This was subtracted from the original signal.

**Figure 3.**
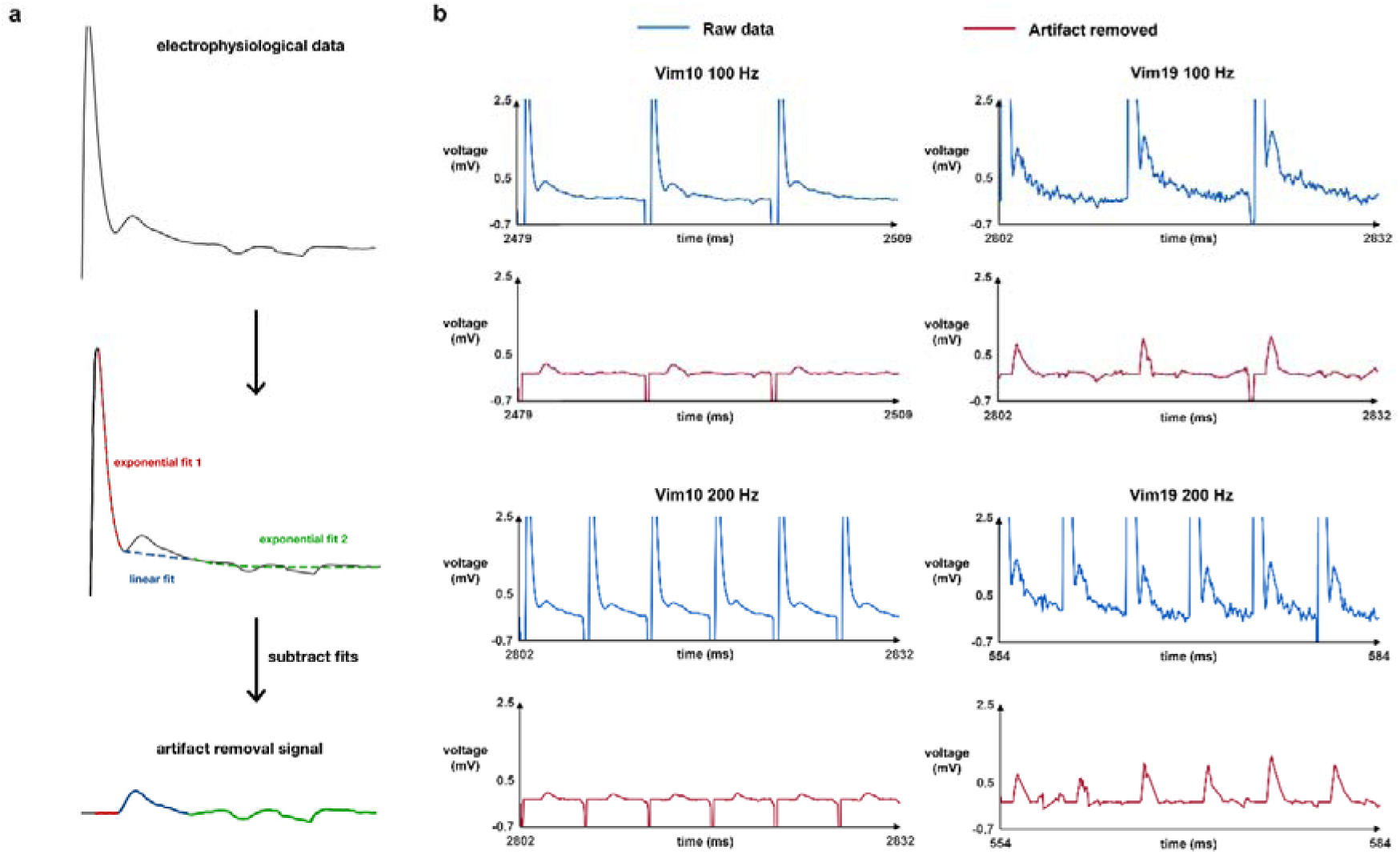
DBS artifact removal preserved quasi-evoked inhibition characteristics: Subfigure **a** is a schematic of the artifact removal methodology used. From the maximum point of the DBS artifact to the starting point of the quasi-evoked inhibition, an exponential fit was applied. Then, from the starting point of the quasi-evoked inhibition to the end, the points were connected using a line. Lastly, from the quasi-evoked inhibition endpoint to the end of the DBS pulse interval, another exponential fit was applied. If no quasi-evoked inhibition was present, a single exponential fit was applied to the entire interval. In subfigure **b**, 30 ms samples are shown from 3 s recordings in two different Vim neurons at both 100 and 200 Hz DBS frequency. The blue recordings represent the unmodified in vivo human extracellular electrophysiological recordings. The pink represents the same time sample after the artifact removal is applied with the preservation of quasi-evoked inhibition.

If no quasi-evoked inhibition was present in an IPI, a single exponential fit was applied to the maximum DBS artifact point to the end of the IPI. This was then subtracted from the raw signal. Furthermore, preprocessing was done to remove spikes, so that exponential fits were not corrupted by the case where an action potential peak was chosen as the endpoint for the fit.

### Detection and fit of quasi-evoked inhibition and firing rate comparison

Once DBS pulse artifacts were removed, quasi-evoked inhibition events were detected using a 0.1 mV threshold on the first 30 ms of each IPI. Next, each event was isolated and received an independent sinusoid fit (Supplementary methods). The individual sinusoid fit parameters were plotted so that their evolution in relation to DBS pulse number could be assessed and correlated with each other in figure 4.

**Figure 4.**
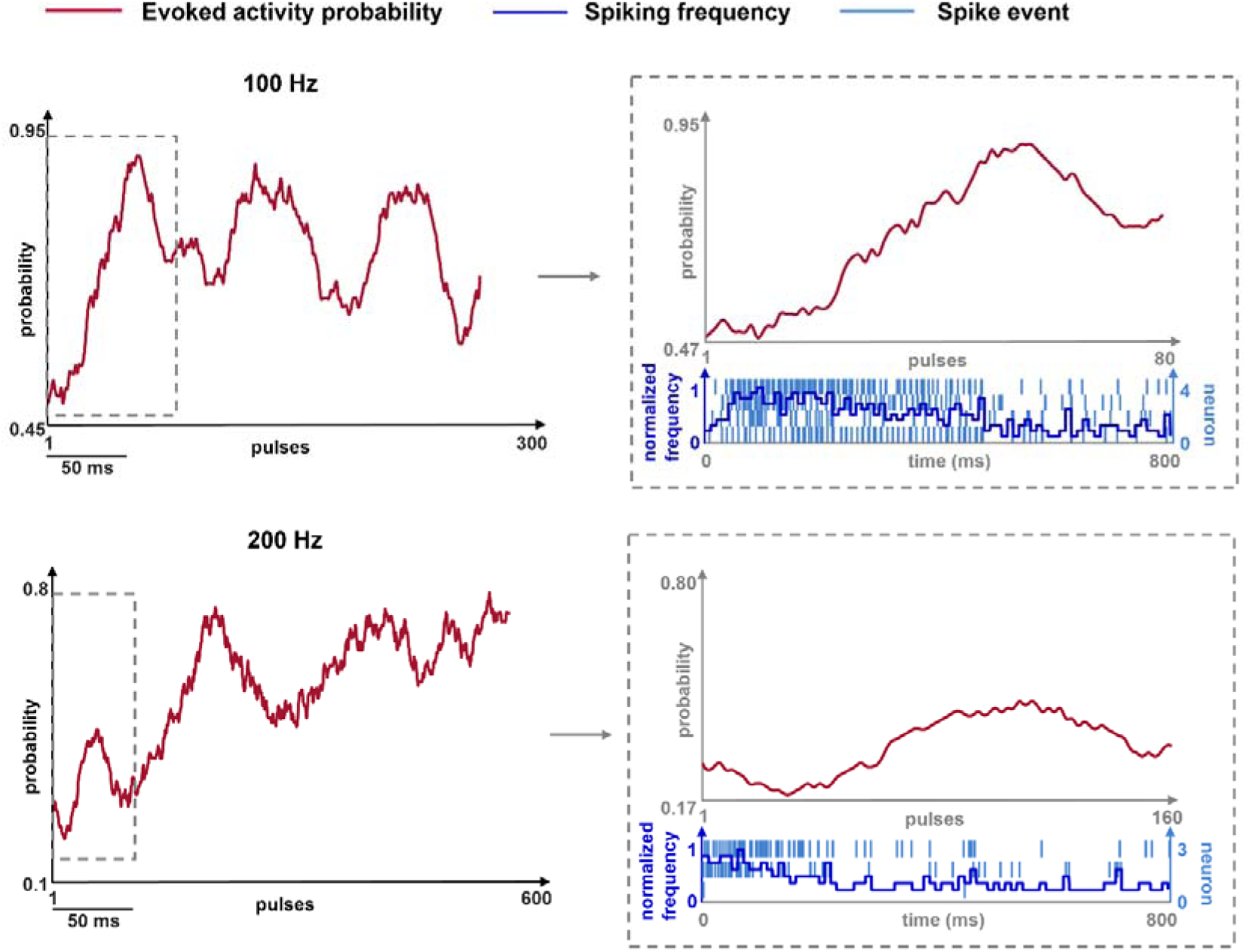
Probability of quasi-evoked inhibition peaks following initial transient excitatory burst: This shows data from all four neurons with quasi-evoked inhibition. For both 100 Hz and 200 Hz, the percentage of neurons with quasi-evoked inhibition following each pulse was computed and then smoothed using a kernel width of 20 pulses for 100 Hz and 35 pulses for 200 Hz (pink). Additionally, the raster plot (light blue) and histogram (dark blue) for the first 800 ms was shown underneath the corresponding probability plot for that time period.

To assess inhibition, quasi-evoked inhibition neurons were separated from neurons without consistent quasi-evoked inhibition. For each frequency, the average steady-state firing rate to baseline firing rate ratio was calculated for each group and then plotted (Supplementary methods). The ratio was taken to control for highly variable baseline firing rates.

To investigate the excitatory burst, raster plots and histograms using 100 bins were plotted for the first 80 ms following the initial DBS pulse. The probability of quasi-evoked inhibition during this time was calculated as a fraction of quasi-evoked inhibition events per DBS pulse and smoothed using a kernel width of 20 pulses for 100 Hz and 35 pulses for 200 Hz (Supplementary methods).

To compare rate of quasi-evoked inhibition and firing rate, the Hideaki method was used to determine optimized kernel sizes for the peri-stimulus time histograms of spiking rate and quasi-evoked inhibition rate (Supplementary methods).^54,55^ This was assessed for individual recordings and then assessed for all 100 Hz recordings and all 200 Hz recordings. Correlations between quasi-evoked inhibition rate and firing rate were computed each time.

### Statistical analysis

To assess the removal of the artifact and preservation of quasi-evoked inhibition signal, the peak amplitude of the quasi-evoked inhibition event in the raw signal and the artifact-removed signal was taken each time. Pearson and distance correlations were taken and a correlation greater or equal to 0.85, accompanied by a significant P-value obtained from a t-test and permutation testing was considered to have strong amplitude preservation. Signal to noise ratios were also taken before and after artifact removal to assess how well the artifact was removed. Additionally, a frequentist and Bayesian right-tailed paired t-tests with unequal variances was used to assess significance. To assess the accuracy of the sinusoid fit, mean squared error (MSE) was computed for each fit.

To statistically compare the firing rate ratios of neurons with quasi-evoked inhibition and neurons without during 100 Hz and 200 Hz stimulation, both one-tailed independent samples t-tests with unequal variances and one-tailed Wilcoxon rank sum tests were conducted and both tests used a 95% confidence interval. Additionally, to compare the state during and after the initial excitatory burst, 20 ratios during the first 60-79 ms to values during 1-20 ms of probability and normalized frequency were computed and a two-tailed Wilcoxon rank sum test with a 95% confidence interval was conducted to determine if there were significant differences between the groups. Lastly, to compare the quasi-evoked inhibition rates to the firing rates, Pearson correlations were taken and a t-test was conducted to determine if correlations were significant. For each Pearson correlation taken, a distance correlation test^56^ was also done with permutation testing to determine significance. This was to validate that computed correlations held true for a test that does not assume linearity or monotonicity.^56^

## Data availability

Single-unit extracellular spiking and field potential available at https://doi.org/10.1101/2020.11.30.404269. Codes used for artifact removal algorithm and analysis are available from ML upon reasonable request.

## Results

### Quasi-evoked inhibition is observed in a subset of Vim neurons following DBS

Quasi-evoked inhibition resulting from DBS was defined as a 2 ms or less positive deflection of the extracellular potential after the peak of the stimulus artifact (Fig. 2, Supplementary Fig. S1). At 100-Hz and 200-Hz DBS, the quasi-evoked inhibition was seen at least once within the first five pulses in each Vim neuron in our 19 neuron dataset (Supplementary Table S1). After its first appearance, in Vim01, Vim04-09, and Vim12-18 the quasi-evoked inhibition halted and never returned, or it returned infrequently for less than five consecutive pulses. In neurons Vim02, Vim10, Vim11, and Vim19 quasi-evoked inhibition halted after pulse five, but returned at approximately pulse 30, after which it frequently returned for at least five pulses. These four neurons were classified as “neurons with consistent quasi-evoked inhibition”, and the other 15 neurons are classified as neurons without consistent quasi-evoked inhibition (pie chart in Fig. 2). Overall, only four out of 19 (or 21.1%) neurons demonstrated this consistent quasi-evoked inhibition in Vim extracellular potential recordings.

### Removal of Vim-DBS artifact with preservation of quasi-evoked inhibition

This DBS artifact removal methodology was used for all four neurons with consistent quasi-evoked inhibition to remove effects from electrode capacitance (Fig. 3, Supplementary Fig. S2)). We observed strong evidence of preservation of quasi-evoked inhibition amplitude features based on Pearson correlation rho measure (median rho = 0.94, 0.89 - 0.95, *P* < 1.00 x 10^-20^). This was confirmed with the distance correlation (median correlation = 0.90, 0.80 - 0.97, *P* < 1.00 x 10^-20^). Additionally, evidence of successful artifact removal was observed in moderately higher signal to noise ratios in the artifact removal signals than the raw data (median_artifact removed_ = 5.1966, median_raw_ = 2.6753, *Z* = 67, *P* = 0.021, BF_10_ = 5.113).

### Probability of quasi-evoked inhibition peaks following initial transient excitatory burst

When evaluating the probability of quasi-evoked inhibition across the entire recording, it ranged from 0.49-0.90 during 100-Hz DBS, and from 0.20-0.78 during 200 Hz DBS (Fig. 4). When assessing the relationship between quasi-evoked inhibition probability and normalized firing frequency, we found strong evidence of highly differing transient state, to post-transient state ratios between the two groups for 100 Hz stimulation (median_transient_ = 0.1667, median_post-transient_ = 1.5349, *Z* = 230, *P* = 3.63 x 10^-8^) and 200 Hz stimulation (median_transient_ = 0, median_post-transient_ = 1.6875, *Z* = 210, *P* = 1.45 x 10^-11^).

### Suppression is stronger following high-frequency Vim-DBS in neurons with quasi-evoked inhibition

We found no statistical difference between the firing rate ratios for 1-10 Hz based on the results of the Wilcoxen rank sum test stimulation (*P*_1 vs 2 Hz_ = 1, *P*_2 vs 3 Hz_ = 0.96, *P*_3 vs 5 Hz_ = 0.72, *P*_5 vs 10 Hz_ = 0.09). However, based on the results of the Wilcoxon rank sum 20-50 Hz stimulation all showed evidence of increased excitation with a significantly higher firing rate ratio compared to 10 Hz stimulation (*Z*_20 Hz_ = 89, *Z*_30 Hz_ = 92, *Z*_50 Hz_ = 96, *P*_20 Hz_ = 0.03, *P*_30 Hz_ = 0.01, *P*_50 Hz_ = 0.002). It is interesting to note that the significance of the p-value and Z score increases with frequency implying stronger an increasing trend in the firing rate ratio from 20- 50 Hz. For 100 Hz, there is a statistically significantly lower firing rate ratio in neurons with consistent quasi-evoked inhibition at 100 Hz (*t*(16) = -3.6409, *d* = -1.7507, *P* = 0.001, BF_10_ = 3.148)). Additionally, the Wilcoxon rank sum test was used to confirm the result for the small sample size (*Z* = 13, *P* = 0.002). It is also interesting to note that the firing rate ratio for neurons without quasi-evoked inhibition during 100 Hz stimulation is greater than one, but less than one for neurons with quasi-evoked inhibition indicating suppression below baseline only in the group with quasi-evoked inhibition (Fig. 1b). For 200 Hz, there is a statistically significantly lower firing rate ratio in neurons with consistent quasi-evoked inhibition at 100 Hz (*t*(8) = -2.8088, *d* = -1.0176, *P* = 0.01, BF_10_ = 2.6839). The Wilcoxon rank sum test does not show significant differences between these two groups due to the extremely small sample size. It is also interesting to note again that the firing rate ratio for neurons without quasi- evoked inhibition during 100 Hz stimulation is greater than one, but less than one for neurons with quasi-evoked inhibition indicating suppression below baseline only in the group with quasi-evoked inhibition (Fig. 1b).

### Quasi-evoked inhibition amplitude follows an oscillatory pattern

As shown in Fig. 5a (Supplementary Fig. S3), the sinusoid amplitude and period were both able to capture the quasi-evoked inhibition intensity and length of occurrence, and the was confirmed by MSE (mean = 2.14 x 10^-2^, 8.54 x 10^-4^ - 5.01 x 10^-2^) demonstrating that our sinusoid model fitted the quasi-evoked inhibition events accurately.

**Figure 5.**
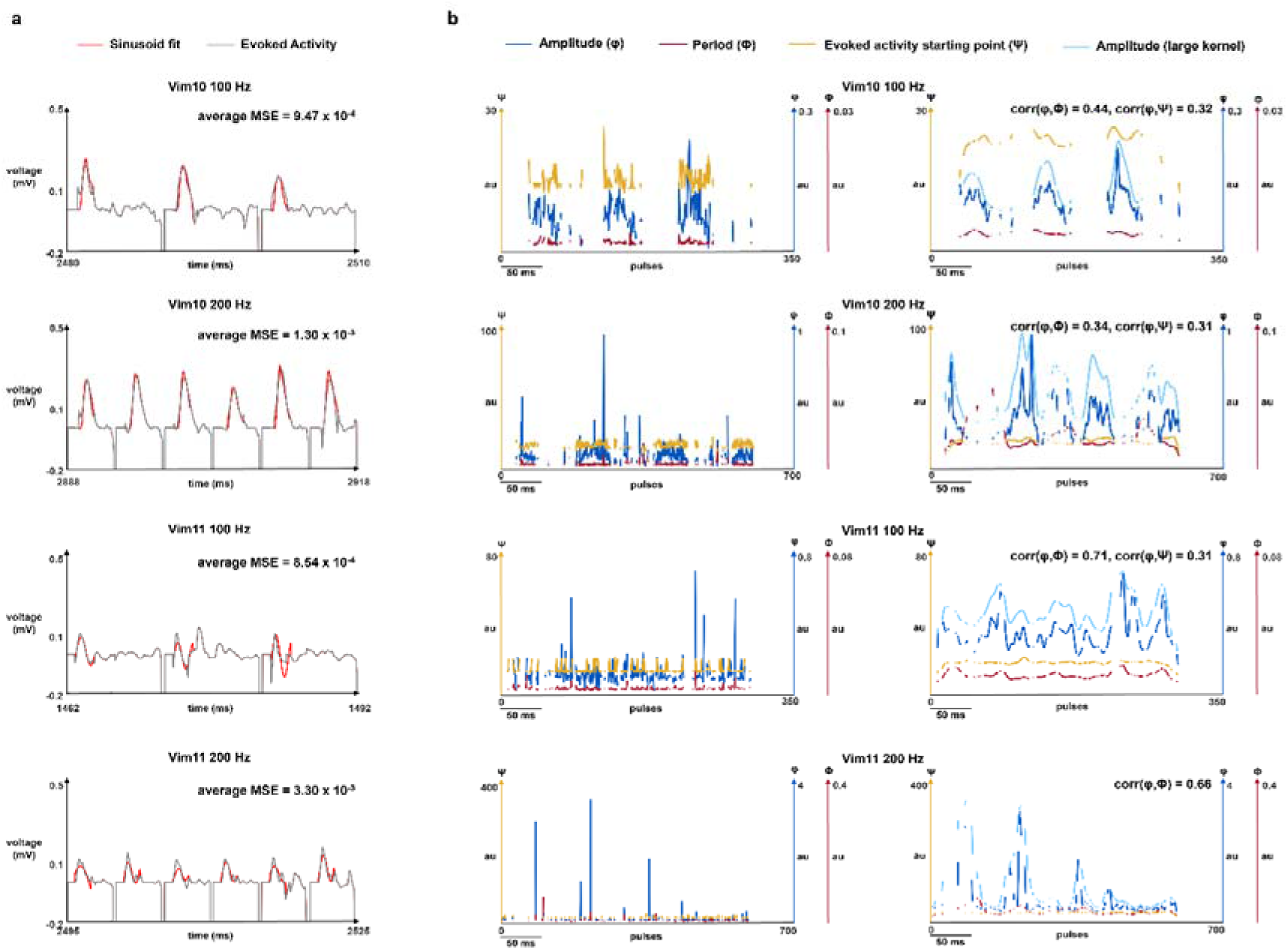
Quasi-evoked inhibition amplitude follows an oscillatory pattern: In subfigure **a**, a sinusoid fit was applied to Vim neuron 10 and Vim neuron 11 at 100 and 200 Hz. The gray recording represents 30 ms samples of 3 s recordings of the in vivo human extracellular electrophysiological recordings with artifact removal applied. The red represents the independent sinusoid fit that was applied to fit each quasi-evoked inhibition event following a DBS pulse. Mean square error over the entire recording is displayed in the top right corner of the plot. Subfigure **b** in the left column shows the parameter values for each sinusoid fit plotted over DBS pulse for Vim10 and Vim11 for 100 Hz and 200 Hz stimulation. The blue curve is the amplitude parameter of the sinusoid fit, the pink curve is the period parameter of the sinusoid fit, and the yellow curve is the starting time point of the quasi- evoked inhibition event within the IPI. If there is no quasi-evoked inhibition following a particular pulse, nothing is plotted and there is a discontinuity in all three curves. Subfigure **b** in the right column shows the smoothed amplitude (light blue), period (pink), and quasi-evoked inhibition start point (yellow) values for each sinusoid fit plotted over DBS pulse for Vim neuron 10 and Vim neuron 11 at 100 and 200 Hz. Plots were smoothed using a Gaussian kernel with widths between four to six. Additionally, the amplitude parameter was also plotted using a Gaussian kernel with width one to two (dark blue). Significant correlations between amplitude and period and correlation between amplitude and quasi-evoked inhibition event start point over the entire recording are both displayed in the top right corner of the plot.

We analyzed three temporal characteristics of the quasi-evoked inhibition events based on the corresponding sinusoid fits: amplitude, period, and the quasi-evoked inhibition event starting time point within the IPI. In figure 5b (Supplementary Fig. S3), we showed how these characteristics evolved in correspondence to each DBS pulse number over the full recordings. The original plots of these characteristics were shown on the left of figure 5b. The smoothed plots (Fig. 5b, right figures) showed oscillatory patterns in the amplitude parameter between 1 - 3 Hz and higher frequency oscillations. Moreover, the amplitude had a significant positive Pearson correlation with the period for each shown recording (median rho = 0.46, 0.15 - 0.71, *P* < 1.00 x 10^-20^) (Fig. 5b, right figures). This was confirmed with the distance correlation (median correlation = 0.43, 0.13 - 0.67, *P* < 1.00 x 10^-20^). The significant positive Pearson correlations between amplitude and quasi-evoked inhibition starting point were in Vim10 100 Hz (rho = 0.32, *P* < 1.00 x 10^-20^), Vim10 200 Hz (rho = 0.31, *P* < 1.00 x 10^-20^), and Vim11 100 Hz (rho = 0.31, *P* < 1.00 x 10^-20^). The corresponding distance correlations were Vim10 100 Hz (correlation = 0.34, *P* < 1.00 x 10^-20^), Vim10 200 Hz (correlation = 0.22, *P* < 1.00 x

10^-20^), and Vim11 100 Hz (correlation = 0.27, *P* < 1.00 x 10^-20^).

### Quasi-evoked inhibition is negatively correlated with the Vim firing rate

We compared the Vim firing rate with the rate of quasi-evoked inhibition in each recording for each neuron receiving HFS (Fig. 6a, Supplementary Fig. S4). We found strong evidence of a significant negative correlation between the quasi-evoked inhibition rate and Vim firing rate, for each recording during 100-Hz stimulation (median rho = -0.24, -0.23 - - 0.77, *P* < 1.00 x 10^-5^, BF_10_ > 300). This was confirmed with the non-signed distance correlation (median correlation = 0.41, 0.22 - 0.79, *P* < 1.00 x 10^-20^). Additionally, we also found strong evidence of a significant negative correlation between the quasi-evoked inhibition rate and Vim firing rate, for each recording during 200-Hz stimulation (median rho = -0.32, -0.26 - -0.51, *P* < 1.00 x 10^-5^, BF_10_ > 300). This was confirmed with the non-signed distance correlation (median correlation = 0.44, 0.29 - 0.55, *P* < 1.00 x 10^-20^). This relationship was also evaluated on the population level and again had strong evidence of negative correlation for 100 Hz stimulation (rho = -0.54, *P* < 1.00 x 10^-5^, BF_10_ > 300) and 200 Hz stimulation (rho = -0.50, *P* < 1.00 x 10^-5^, BF_10_ > 300), and confirmed with non-signed distance correlation for 100 Hz stimulation (correlation = 0.57, *P* < 1.00 x 10^-20^), and 200 Hz stimulation (correlation = 0.49, *P* < 1.00 x 10^-20^).

**Figure 6.**
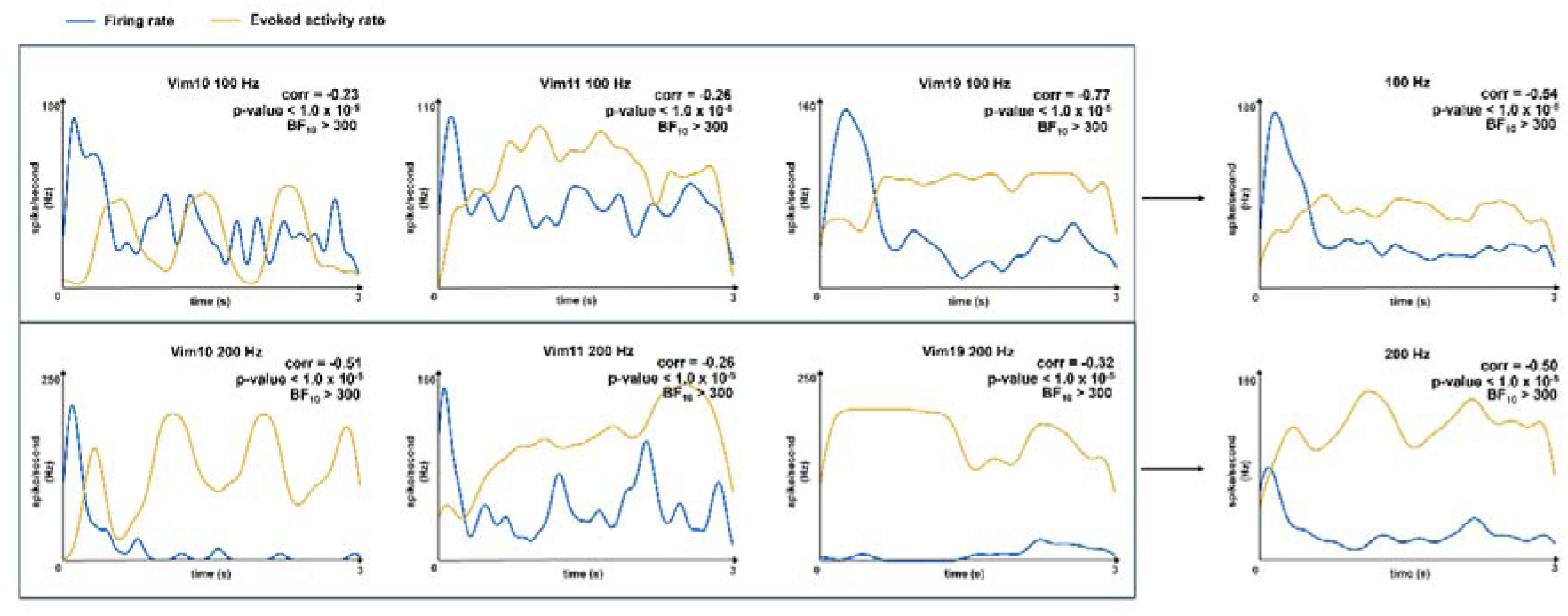
Quasi-evoked inhibition is negatively correlated with the Vim firing rate: In subfigure **a**, all four neurons that exhibit quasi-evoked inhibition are shown over their full electrophysiological recordings ranging from 3.0-5.0 s. The rate of presence of quasi-evoked inhibition is computed using an optimized width of a Gaussian kernel. This is plotted in yellow over the entire time interval. Firing rate is also computed using a width consistent with the Hideaki optimized Gaussian kernel method. This is plotted in blue over the entire time interval. Correlation and P-values are computed between the two curves and displayed in the top right of each plot. Subfigure **b** shows data from all four neurons with quasi-evoked inhibition.

## Discussion

In this work, we conducted an in-depth analysis of a positive-going evoked potential seen in a subset of neurons during HFS of the Vim and evaluated evidence of its possible mechanism in network inhibition. Our results show evidence of synaptic depression and network inhibition recruitment during HFS of the Vim. Previous research has questioned whether DBS enacts inhibition on the cellular level^29,30,57–59^ or through network effects^33,60–62^, and here we demonstrate that at high frequencies, there is support for both proposed DBS mechanisms contributing towards inhibition.

We utilized microelectrode recordings during DBS that were able to track neuronal signals with high spatial resolution on the single-unit level, so that quasi-evoked inhibition could be clearly identified and temporally analyzed. In contrast, the local field potential data recorded with DBS macroelectrodes lack the tractability of single-unit level neuronal activities.^63–65^ Other studies have benefitted from the use of microelectrode recordings in the Vim. Milosevic et al., (2018) temporally analyzed accelerometry recordings in relation to Vim bursting and suppression,^28^ and Kon Kam King et al. (2017) classified movement-responsive cells and analyzed single-unit features like firing rate and bursting.^66^ However, to the best of our knowledge, this is the first work to identify a positive-going evoked potential in Vim single-unit recordings and investigate it in relation to temporal characteristics like instantaneous firing rate.

In most of the HFS recordings, there was a transient period of excitation followed by a suppressed steady-state response. This is consistent with previous literature characterizing Vim firing during HFS.^28,66,68^ However, this is inconsistent with other DBS target nuclei, like the STN and the internal globus pallidus that are suppressed during HFS without an initial period of excitation.^69–71^ Milosevic et al. (2018) discusses various possible mechanisms for this response in the Vim^28^; then, hypothesizes that it is the result of the the high prevalence of glutamatergic inputs^45,67^ activated by DBS leading to either vesicle depletion or presynaptic calcium reduction, both result in synaptic depression.^28–30,72–74^ This is consistent with the results seen in the frequency-dependent suppression experiments conducted in figure 1. At very low frequency DBS, the firing is only slightly above baseline which may be because there is not frequent enough stimulation to strongly activate the glutamatergic terminals. Then, from 20-50 Hz stimulation, the firing rate increases from baseline as the stimulation frequency increases. This could be due to the activation of the glutamatergic terminals, but not at a high enough frequency to induce vesicle depletion or calcium reduction. Furthermore, we found that quasi-evoked inhibition does not emerge until after the initial excitatory burst as its probability is at a global minimum during the transient burst (Fig. 4). Additionally, quasi- evoked inhibition probability reaches a local maxima right after the transient burst (Fig. 4). Consequently, the HFS excitatory burst could be what allows for the recruitment engagement of inhibitory neurons. This hypothesis is consistent with thalamo-cortical circuit conduction delays. The conduction delay from thalamus to cortex is estimated to be about 8 ms,^75^ and within cortex transmission delays are 10 ms.^76,77^ Also, Huo et al. (2020) found that using a cortical to thalamic conduction delay of about 3 ms,^78^ inactivation of the cortex only took about 7.3 ms to reduce thalamic activity in areas investigated, including the TRN.^79^ Lastly, the synaptic transmission delay within thalamus is estimated as 1 ms.^80^ The first local maxima in quasi-evoked inhibition probability is at about 55 pulses for 100 and 200 Hz stimulation (Fig. 4) which corresponds to about 55 ms and 27.5 ms. Even with all of the transmission delays discussed, it is physiologically feasible for glutamatergic signals to be sent from the Vim to cortex to the TRN, and then GABAergic signalling to be received by the Vim before first maxima in quasi-evoked inhibition probability.

Next, we found that steady state firing rate controlled for baseline firing rate is at its lowest during Vim HFS in all neurons, which supports that HFS causes cellular synaptic depression leading to reduced firing (Fig. 1b). As mentioned previously, this could be due to DBS activating the glutamatergic terminals at such a high frequency that it leads to presynaptic calcium reduction or vesicle depletion.^28,72–74^ However, even with the steady-state firing rate at its lowest compared to baseline during HFS, the neurons with quasi-evoked inhibition were the only ones that experienced suppression below baseline during HFS (Fig. 1b, blue samples), and this was significantly lower than the group of neurons without quasi- evoked inhibition. During DBS, evoked potentials have been found both at the stimulation site,^81–84^ and at cortical efferents.^81,85,86^ Many evoked potentials observed at the stimulation site are believed to be a result of synchronous activation of local neurons.^81,82^ However, there is support for the hypothesis that the evoked resonance neural activity observed during STN-DBS is partially due to reciprocal inhibition from the GPe.^34,84^ Previous modelling of Vim-DBS shows that neuronal suppression and inhibition engagement can only be achieved during HFS,^37^ which is supported by increased steady state firing compared to baseline observed during lower frequency stimulation. Consequently, this supports the claim that these neurons that display quasi-evoked inhibition are suppressed beyond the means of synaptic depression and also by network reciprocal inhibition engagement from the TRN and thalamic interneurons.

In addition, we showed that there are oscillations in the amplitude parameter of the quasi-evoked inhibition fit in every recording (Fig. 5). Previous research has shown high frequency oscillations of evoked resonance neural activity in the STN.^81,84,87^ While there is no current evidence that quasi-evoked inhibition is resonant following the cessation of DBS, the amplitude oscillating at a frequency other than the stimulation frequency or the frequency of cardioballistic artifacts may suggest a neural causation. Modelling has shown that network inhibition response during Vim HFS oscillates.^37^ Our results support this because the amplitude increase can be a result of a facilitatory plasticity effect on the quasi-evoked inhibition from increased TRN and interneuron inhibitory inputs. Then, the amplitude decrease can be a result of a depression effect on the quasi-evoked inhibition from decreased TRN and interneuron inhibitory inputs. Since this repeats throughout the recording, it supports the idea of cyclical inhibitory inputs recruited by Vim HFS. Moreover, the higher frequency oscillations in the amplitude parameter are qualitatively at a higher frequency for 200 Hz than 100 Hz in each sample (Fig. 5). This implies that different stimulation frequencies may modulate the amplitude differently. Additionally, the overall scalar amplitude value was qualitatively higher during 200 Hz stimulation compared to 100 Hz stimulation (Fig. 5). Both of these findings are consistent with literature involving the STN. Ozturk et al. (2021) found the amplitude of the evoked potentials observed were significantly higher during clinically effective stimulation frequencies;^87^ while stimulating above 100 Hz is effective in the Vim,^88^ 200 Hz microstimulation is more clinically effective.^28^ Furthermore, the lowest overall amplitude was observed in Vim10 and Vim11 at 100 Hz (Fig. 5). These neurons also experienced some of the highest steady-state firing rates in the quasi-evoked inhibition group (Fig. 6). While more testing needs to be done to confirm findings, this may suggest that overall amplitude of quasi-evoked inhibition is higher with stronger inhibitory inputs, adding more support to the plasticity mechanism of quasi-evoked inhibition oscillations suggested.

Lastly, we showed that quasi-evoked inhibition and firing rate are significantly anticorrelated within individual HFS recordings (Fig. 6). Also, the quasi-evoked inhibition rate for 200 Hz had a higher overall frequency than 100 Hz (Fig. 6). Since 200 Hz microstimulation is more clinically effective and results in higher levels of cell inhibition,^28^ this might suggest a mechanism for that clinical observation. If quasi-evoked inhibition is a temporal marker of inhibitory inputs and it appears at a much higher frequency during stimulation that results in higher cell inhibition, it could mean that 200 Hz microstimulation allows for optimal engagement of inhibitory inputs, leading to the observed firing suppression.^28^ Additionally, this supports the idea that quasi-evoked inhibition could be a temporal marker of when individual neurons are experiencing network inhibition and are not just being suppressed by cellular depression. Having a potential biomarker of when network inhibition is engaged could be utilized during DBS electrode placement procedures, as microelectrode recordings are commonly used during the procedure.^89–91^ While clinical correlations are yet to be investigated, at the stimulation target, quasi-evoked inhibition may be observed at higher frequencies as it results in higher cell inhibition (Fig. 1, Fig. 6). Also, quasi-evoked inhibition could be used as an additional clinical recording to improve the stimulation parameters of Vim-DBS. Vim-DBS parameter optimization can improve clinical outcomes,^92,93^ and having more data about the thalamo-cortical network could lead to better data fitting for optimization strategies. Moreover, a temporal biomarker of inhibitory projections to the Vim could allow for more insightful investigation into if inhibition engagement affects movement or cognitive responses.

### Limitations and future directions

In this work, quasi-evoked inhibition was found to be associated with increased neuronal suppression both overall and temporally. However, firing rate is only weakly anticorrelated with tremor suppression.^28^ We plan to assess stronger clinical correlates, like accelerometer and EMG recordings, with quasi-evoked inhibition in future works.

Furthermore, “tremor cells” are characterized by periodic oscillation in rate of discharge synchronous with limb tremor and found in essential tremor patients.^66,94,95^ While additional characteristics of these cells have not been clearly identified in the present study, Lenz et al. (1994) noted that they are unresponsive to sensory stimulation.^95^ These intrinsic attributes of “tremor cells” are critical to understanding the pathology of essential tremor, but investigating this in relation to the network is also critical. The quasi-evoked inhibition classified in this work may be a biomarker of thalamocortical network engagement, so that in future works tremor cells can be investigated on a network level.

Another limitation of this work is that ballistocardiogram artifacts were not considered. Ballistocardiogram artifacts have been detected^96^ and removal has been attempted^97–100^ in EEG recordings. Any cardiac related artifacts were not detected nor removed in this work. However, since the sinusoid parameters, especially amplitude, smoothed with a large kernel, oscillated at similar frequencies to cardiac activity, it is possible that these oscillations were a result of a ballistocardiogram artifact. For this reason, no possible neural mechanistic causes or conclusions about these slow oscillations were discussed, as an obvious confounding variable may be affecting results.

Additionally, many DBS targets have been identified with the use of MRI and other medical imaging technologies.^101–105^ We hypothesize that the best DBS target would involve neurons with tremor related activity that can engage with the thalamocortical network. In the future, our work will be assessed in relation to imaging modalities. Other limitations include a small sample size, and while there is experimental evidence and corresponding modelling^37^ to suggest inhibitory inputs, we do not have corresponding neuron recordings to evaluate this definitively.

## Conclusion

A subset of Vim neurons were found to be suppressed beyond the level of cellular depression during HFS. These results support both synaptic depression and network inhibition hypotheses of DBS mechanisms. Moreover, HFS of the Vim may cause antidromic and orthodromic activation of cortical motor areas, which in turn excite the TRN, the inhibitory projection to the Vim. The transient burst exhibited by the Vim in response to HFS might be what allows for the recruitment of inhibitory populations. The Vim neurons suppressed significantly more than the controls exhibit quasi-evoked inhibition which may be a biomarker of when inhibitory populations are engaged and projecting GABAergic signals reciprocally. These findings provide new insight into the network mechanism underlying Vim-DBS. In turn, this may provide new methodologies for microelectrode guided essential tremor stimulation protocols, targeting, and DBS parameter optimization.

## Acknowledgements

We would like to thank the patients that participated in this work.

