## Supplementary for "Frequency-dependent Inhibition during Deep Brain Stimulation of Thalamic Ventral Intermediate Nuclei"

**Supplementary methods**

**Artifact removal**

Exponential fit methodologies have been developed to remove deep brain stimulation (DBS) artifacts.^1^ However, our goal was to remove the DBS artifact without modifying the quasi-quasi-evoked inhibition markers which were often merged with the artifact. Consequently, the DBS artifact removal was completed in three distinct parts to preserve the features of the quasi-quasi-evoked inhibition. The first step was to apply an exponential fit up until the start point of the quasi-quasi-evoked inhibition. Then, a linear subtraction was applied to the section of the interpulse interval (IPI) containing the quasi-quasi-evoked inhibition, so that the electrophysiological signal was not altered in the process. Lastly, a second, separate, exponential fit was applied following the quasi-quasi-evoked inhibition for the remainder of the IPI.

(1) In the first, exponential fit step of the artifact removal process, we localized the time point of the maximum voltage of the stimulus artifact. The first exponential fit was applied starting 0.48 ms after this time point and anything prior to that in the IPI was set to 0. To find the endpoint of the first exponential fit, the MATLAB “findpeaks” function was used on the negative signal to find local minima. If local minima were detected, the endpoint for the exponential fit was set to 0.08 ms prior to the first local minima. Using the MATLAB fit function, an exponential fit using parameters A, g, and O, and equation:

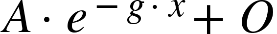

The initial parameters used were A = 50, g = 0, O = 5000 was applied to the electrophysiological data between the determined start and endpoints. Once the fit was completed, it was subtracted from the original electrophysiological data between the start and endpoints.

(2) The linear subtraction step of the artifact removal process was only completed if there was quasi-quasi-evoked inhibition within the IPI (Fig. 3a). The starting point was set to the first local minima. Through visual inspection of the electrophysiological data, quasi-quasi-evoked inhibition did not last more than 2 ms. For this reason, the linear subtraction endpoint was set to the time point 2 ms following the start point. The start and endpoints were connected using a line. This was then subtracted from the original electrophysiological data between the start and endpoints.

(3) The second exponential fit step of the artifact removal process was only completed if there was quasi-quasi-evoked inhibition within the IPI. The start point was set to 0.08 ms following the linear subtraction endpoint. The endpoint was set to the time point 0.40 ms before the end of the IPI. Using the MATLAB fit function, an exponential fit using parameters A, g, and O, and equation:

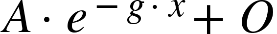

The initial parameters used were A = 5, g = 10, O = 500 was applied to the electrophysiological data between the determined start and endpoints. Once the fit was completed, it was subtracted from the original electrophysiological data between the start and endpoints (Fig. 3a).

As shown in the schematic (Fig. 3a), the DBS pulse artifact and the rest of the inter-pulse recording followed a clear exponential shape; this resulted in minimal error in the exponential fits. During the artifact removal, the quasi-quasi-evoked inhibition was preserved by a linear fit (Fig. 3a), which prevents the quasi-quasi-evoked inhibition from being corrupted by the curvature of a nonlinear fit.

In rare cases, a spike depolarization peak was chosen as a fit start or endpoint. This resulted in an inaccurate fit that always undershot the electrophysiological data and artifact removal signals that were too high. To correct this, before applying artifact removal, we removed the spikes that could alter the fit. We found any instances in the raw data of continuous increase of 0.12 mV or more from a local minima, excluding DBS pulses. This was classified as a spike. Then, the electrophysiological data was modified to introduce a line connecting the point 0.4 ms before and 0.4 ms after the local minima. The spike time was found using the MATLAB “findpeaks” function on the negative signal. Any potential spike within the first 1.6 ms of the IPI was left unmodified to prevent the modification of any quasi-quasi-evoked inhibition. Furthermore, if no quasi-quasi-evoked inhibition could be detected within the IPI, then a single exponential fit was applied to the entire IPI excluding the first 0.48 ms and last 0.40 ms, which may include artifacts from the following DBS pulse.

**Quasi-quasi-evoked inhibition detection**

Once artifact removal was completed, quasi-quasi-evoked inhibition events could be detected using simple thresholding. To exclude any low amplitude quasi-quasi-evoked inhibition events, a threshold of 0.1 mV was used on the artifact removed signal. Low amplitude quasi-quasi-evoked inhibition events were excluded because they were sometimes observed at 30-50 Hz DBS, and high amplitude events were unique to high-frequency stimulation. Various thresholds between the range 0.75 - 1.25 mV were tested but resulted in little change to the correlation between quasi-quasi-evoked inhibition rate and spiking rate. Consequently 0.1 mV was used as it best reflected the quasi-quasi-evoked inhibition amplitude of all neurons. The threshold was applied in the MATLAB "findpeaks" function to find the time points of quasi-quasi-evoked inhibition maxima.

**Sinusoid fit of quasi-quasi-evoked inhibition**

We applied a sinusoid fit to characterize the temporal behaviour of the quasi-quasi-evoked inhibition throughout the entire recording. The start point of the fit was set to the time point of the local minima that was immediately prior to the quasi-quasi-evoked inhibition maxima. The quasi-quasi-evoked inhibition maxima and minima were found using the MATLAB “findpeaks” function on both the positive and negative artifact removal signal. If none could be found, the latest time point set to zero at the beginning of the IPI was used as the start point. The endpoint was set to the time point 2 ms after the start point. Using the MATLAB fit function, a sinusoid fit using parameters A, B, C, and D, and equation:

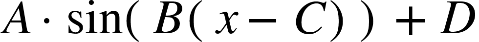

The initial parameters used were A = 100000, B = 100, C = 5, D = 1 was applied to the electrophysiological data processed by artifact removal between the determined start and endpoints. Next, we computed the mean squared error (MSE) between the artifact removed signal and the sinusoid fit. If the MSE > 0.01 or the maxima of the artifact removed signal and the sinusoid fit were not within 0.05 mV of each other, the fit was determined to be inaccurate and modified by the following method. The same start and endpoints were used. Next, Using the MATLAB fit function, a sinusoid fit using parameters A, B, C, and D, and equation:

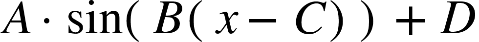

The initial parameters used were random within ranges 1 <= A <= 10000, 1 <= B <= 10000, 1 <= C <= 100, 1 <= D <= 10 was applied to the artifact removed signal. The MSE between the artifact removed signal and the sinusoid fit was again computed. This process was repeated up to 50 times. The MSE of each acceptable fit was saved, and for each neuron with quasi-quasi-evoked inhibition, a simple average of the MSEs was taken and reported. If there were any obvious inaccuracies in the sinusoid fit, they were manually computed afterwards. Each quasi-quasi-evoked inhibition was fit independently using this process.

The period was computed from the B parameter and the amplitude, period, and quasi-quasi-evoked inhibition starting time point were plotted for each DBS pulse number. The amplitude, period, and quasi-quasi-evoked inhibition start point curves were smoothed using a Gaussian kernel with kernel sizes ranging from four to six DBS pulses. Once a smooth plot was constructed, Pearson and distance correlations were computed between amplitude and period, and between amplitude and quasi-quasi-evoked inhibition starting time point. In the computations, we removed the IPIs where there were no quasi-quasi-evoked inhibition events, because in this case no fit was applied.

**Inhibition quantification**

To quantify suppression, the ratio between the DBS average steady state firing rate and the baseline average firing rate was taken. Since each neuron had recordings during at least two DBS frequencies, the electrophysiological data prior to the first DBS pulse from all recordings of the same neuron were concatenated to form the baseline. For 1 Hz DBS, the last two pulses that included the whole 1 s IPI were considered. For 2-5 Hz DBS, the last three pulses that included the full IPI were considered. For 10 Hz, the last five pulses that included the full IPI were considered. For 20-50 Hz, the last 15 pulses that included the full IPI were considered. Lastly, for 100-200 Hz, the last 10 pulses that included the full IPI were considered. The number of IPIs were chosen to ensure the exclusion of the transient state and to keep the total time interval being evaluated relatively consistent between frequencies. For each neuron, the average baseline firing rate and average firing rate at each available frequency (for pulses previously described) were computed and then the ratio between DBS steady state firing and baseline firing at each frequency was taken. For frequencies 1-50 Hz only ratios for no quasi-quasi-evoked inhibition activity were plotted because only Vim02 had available low frequency recordings in the quasi-quasi-evoked inhibition group (Supplementary Table. S1). For each frequency, all available firing rate ratios were averaged, and the standard deviation was calculated. Both the average and the standard deviation were plotted for each frequency. Next, in 100-200 Hz frequencies, the neurons were separated into quasi-quasi-evoked inhibition and no quasi-quasi-evoked inhibition groups, so that the no quasi-quasi-evoked inhibition group could be used as a control. The 100 Hz quasi-quasi-evoked inhibition group included all four neurons with quasi-quasi-evoked inhibition and the 100 Hz no quasi-quasi-evoked inhibition group included the 14 remaining available recordings. In both groups, the average firing rate ratio was computed along with the standard deviation for each stimulation frequency and these were plotted. Moreover, a Bayesian left-tailed independent samples t-test with unequal variances and a left-tailed Wilcoxon rank sum test was conducted. The 200 Hz quasi-quasi-evoked inhibition group included Vim10, Vim11, and Vim19. The 200 Hz no quasi-quasi-evoked inhibition group included the rest of the available 200 Hz stimulation Vim recordings. Again, in both groups, the average firing rate ratio was computed along with the standard deviation and these were plotted. A second Bayesian left-tailed independent samples t-test with unequal variances and left-tailed Wilcoxon rank sum test was conducted for the 200 Hz groups.

**Comparison between quasi-quasi-evoked inhibition and spike rate**

To compare the rate of quasi-quasi-evoked inhibition and spiking, a Gaussian kernel was used. The Hideaki method^2,3^ was first applied to the saved binary spike trains for the full electrophysiological recording of each neuron with quasi-quasi-evoked inhibition during 100-Hz and 200-Hz high frequency stimulation to determine an optimized kernel size for the spike rate. Using the saved time points of the maxima of each quasi-quasi-evoked inhibition event, the same Hideaki method was applied to the full electrophysiological recording of each neuron with quasi-quasi-evoked inhibition during 100-Hz and 200-Hz DBS to determine an optimized kernel size for rate of quasi-quasi-evoked inhibition. Vim02 was only stimulated at 100 Hz, so 200 Hz data was not available (Supplementary Table S1). The kernel sizes were modified so that they would become more consistent with each other while remaining consistent with the optimized kernel sizes. The Pearson correlation and associated p-value and bayes factor between quasi-quasi-evoked inhibition rate and spiking rate were then calculated and reported. Since the distance correlation could not be computed with a sample size greater than 5000, 5000 evenly spaced samples were taken across the entire recording to compute the distance correlation. Permutation testing was done to determine each distance correlation significance.

Next, we compared the percentage of quasi-quasi-evoked inhibition between all four recordings that displayed quasi-quasi-evoked inhibition. For both 100 Hz and 200 Hz, out of all recordings with quasi-quasi-evoked inhibition, the smallest total number of DBS pulses in a recording was considered. For each pulse, a percentage of neurons exhibiting quasi-quasi-evoked inhibition following the pulse was computed (Fig. 5b). Then, the probability was computed as the average value across 20 following pulses at 100-Hz DBS, and across 35 following pulses at 200-Hz DBS. Due to this average computation, the curve ends at 281 pulses instead of 300 pulses for 100 Hz and ends at 566 pulses instead of 600 for 200 Hz. This was then plotted using the MATLAB “interp1” function. For the first 800 ms, the raster plots were constructed for neurons with quasi-quasi-evoked inhibition at 100 Hz and 200 Hz. Also, a histogram was constructed with 8 ms bins to assess the group firing frequency. To assess whether the quasi-quasi-evoked inhibition probability peaked after the excitatory burst, 20 separate ratios during the first 60-79 ms to values during 1-20 ms of probability and normalized frequency were computed. Next a two-tailed Wilcoxon rank sum test was taken to assess statistical significance.

**Supplementary figures**

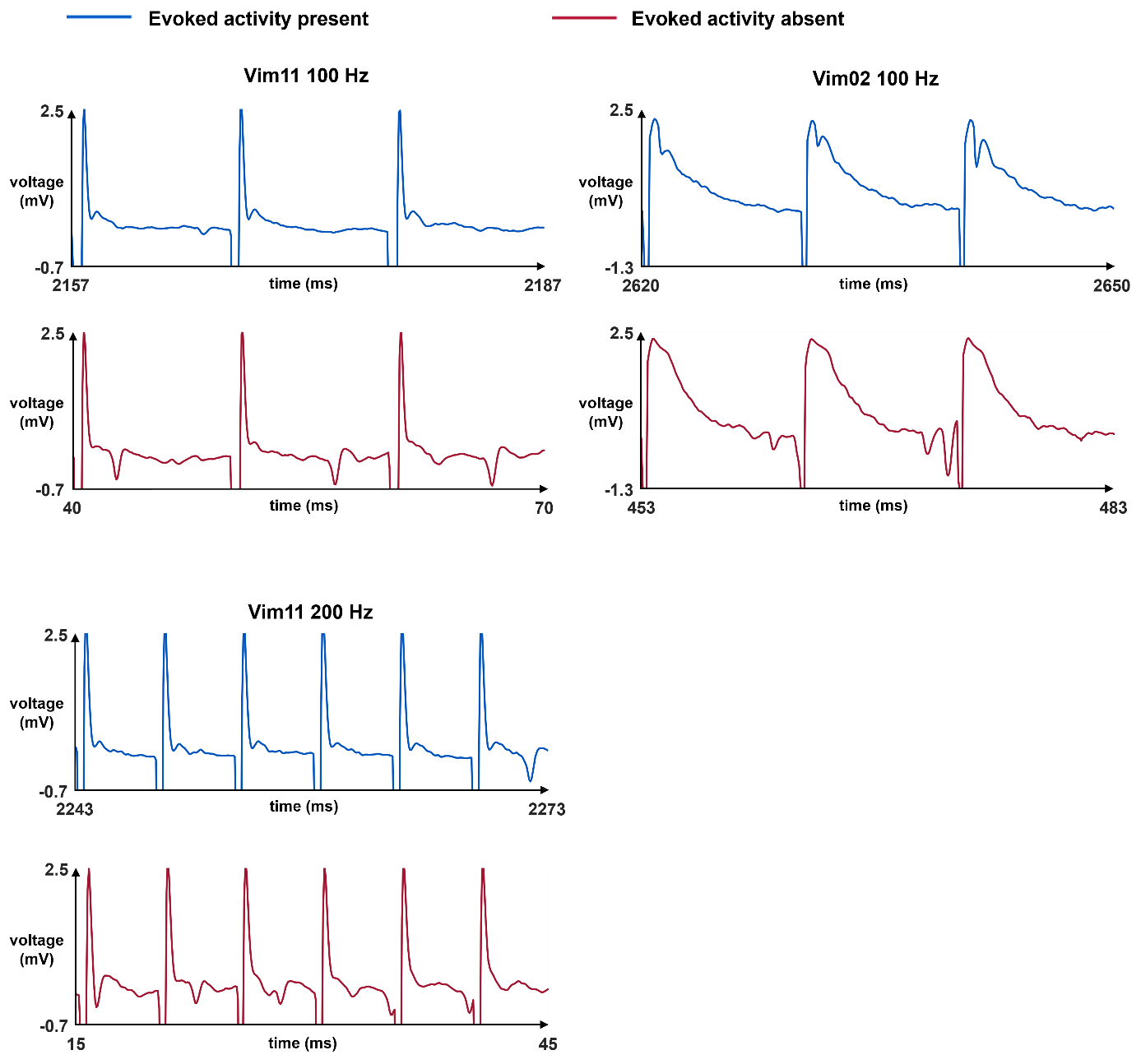

**Supplementary Figure S1 Presence of quasi-quasi-evoked inhibition in Vim-DBS electrophysiological data:** In vivo human extracellular electrophysiological recordings are measured over time. 30 ms samples are shown from 3 s recordings in two different Vim neurons at both 100 and 200 Hz DBS frequency. The blue recordings represent areas of the recording with quasi-quasi-evoked inhibition present following the DBS pulse. The pink recordings represent another area of the same recording where no quasi-quasi-evoked inhibition following the DBS pulse can be observed.

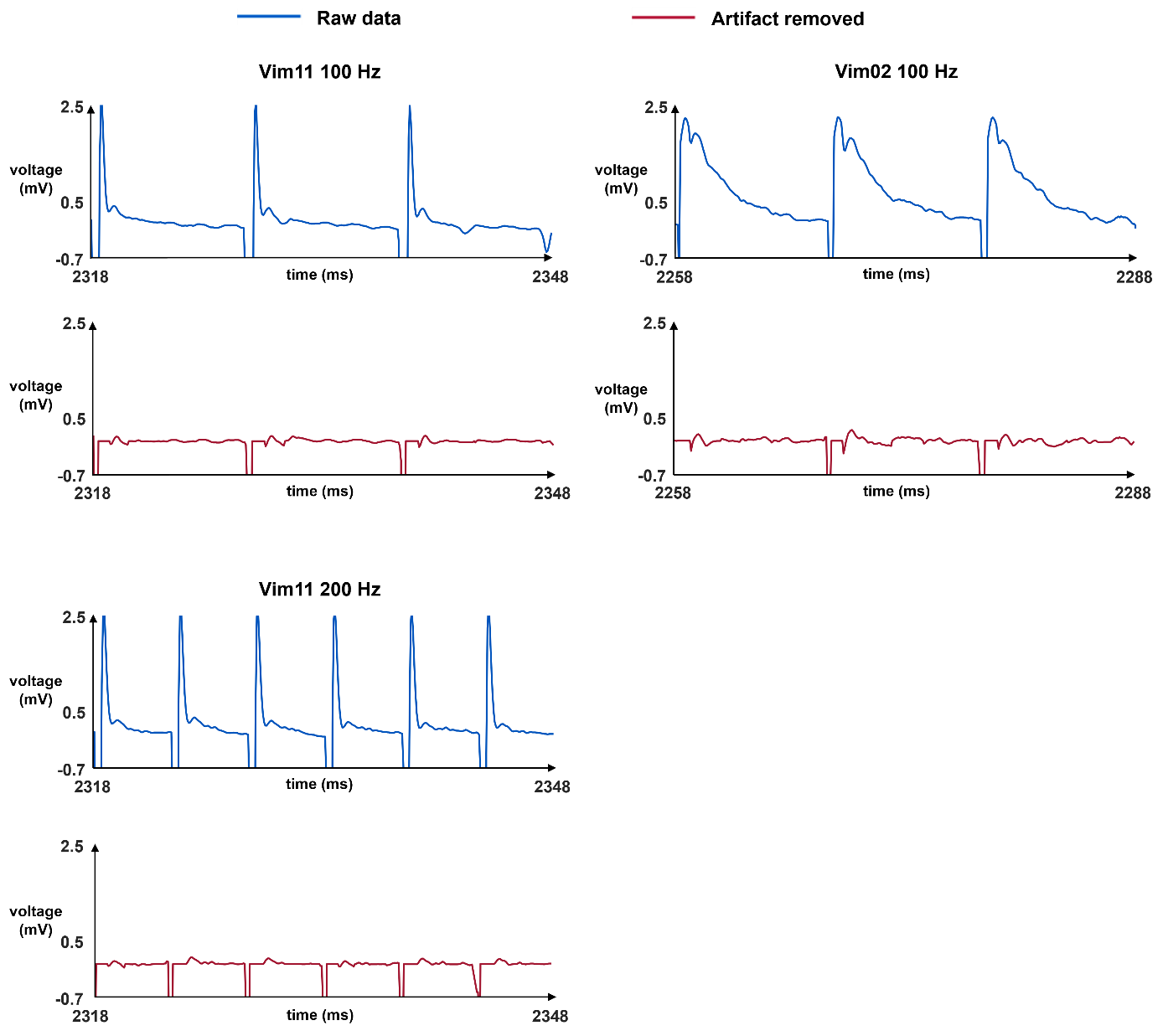

**Supplementary Figure S2 Removal of Vim-DBS artifact with preservation of quasi-quasi-evoked inhibition:** 30 ms samples are shown from 3 s recordings in two different Vim neurons at both 100 and 200 Hz DBS frequency. The blue recordings represent the unmodified in vivo human extracellular electrophysiological recordings. The pink represents the same time sample after the artifact removal is applied with the preservation of quasi-quasi-evoked inhibition.

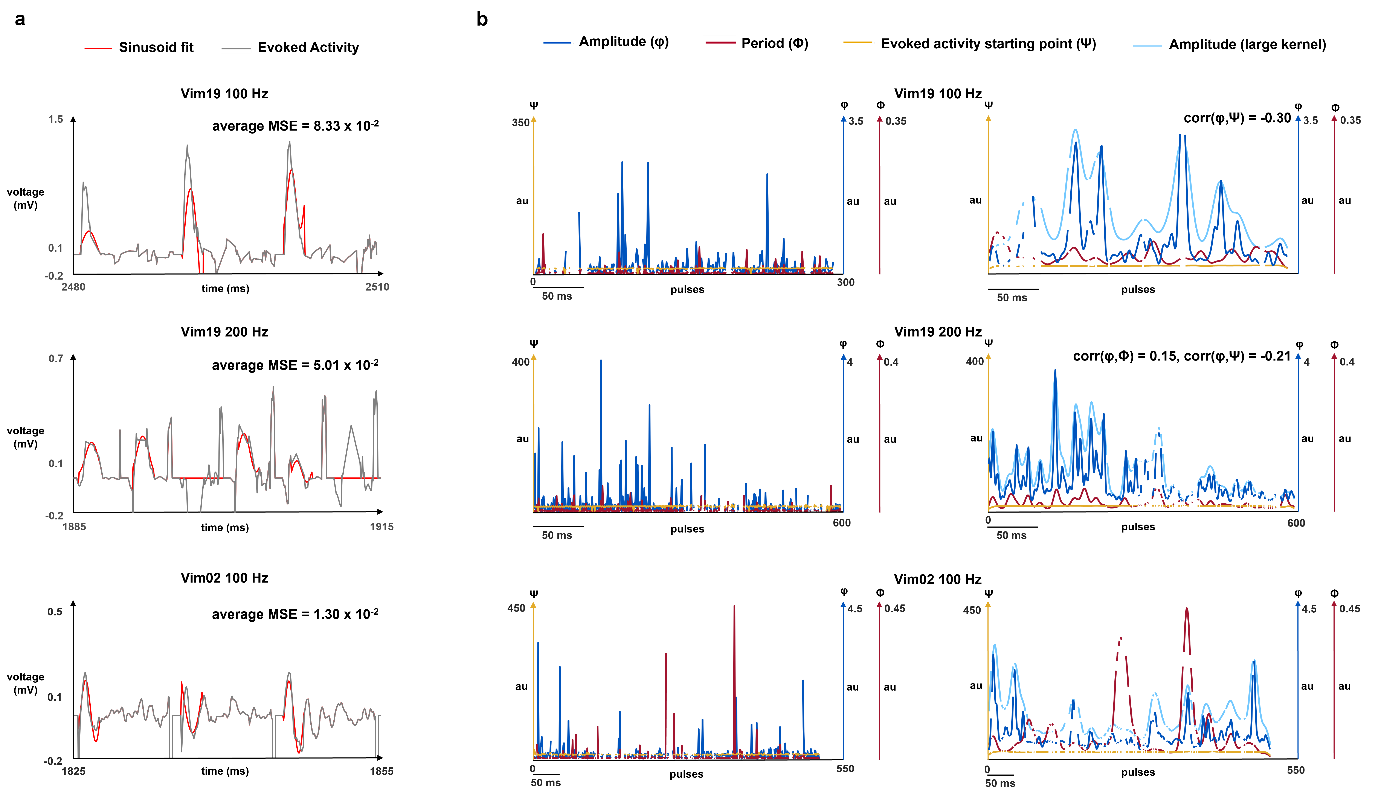

**Supplementary Figure S3 Sinusoid fit of Vim-DBS quasi-quasi-evoked inhibition and parameter evolution:** In subfigure **a**, a sinusoid fit was applied to Vim neuron 19 at 200 Hz and Vim neuron 02 and Vim neuron 19 at 100 Hz. The gray recording represents 30 ms samples of 3 s recordings of the in vivo human extracellular electrophysiological recordings with artifact removal applied. The red represents the independent sinusoid fit that was applied to fit each quasi-quasi-evoked inhibition following a DBS pulse. Average mean square error over the entire recording is displayed in the top right corner of the plot. Subfigure **b** in the left column shows the parameter values for each sinusoid fit plotted over DBS pulse for Vim neuron 19 at 200 Hz and Vim neuron 02 and Vim neuron 19 at 100 Hz. The blue curve is the amplitude parameter of the sinusoid fit, the pink curve is the period parameter of the sinusoid fit, and the yellow curve is the starting time point of the quasi-quasi-evoked inhibition within the inter-DBS pulse interval. If there is no quasi-quasi-evoked inhibition following a particular pulse, nothing is plotted and there is a discontinuity in all three curves. Subfigure **b** in the right column shows the smoothed parameter values for each sinusoid fit plotted over DBS pulse for Vim neuron 19 at 200 Hz and Vim neuron 02 and Vim neuron 19 at 100 Hz. Plots were smoothed using a Gaussian kernel with widths between four to six for amplitude (light blue), period (pink), and quasi-quasi-evoked inhibition start point (yellow). Additionally, amplitude was plotted again (dark blue) using a Gaussian kernel sized one to two. Correlation between amplitude and period and correlation between amplitude and quasi-quasi-evoked inhibition start point over the entire recording are displayed in the top right corner of the plot.

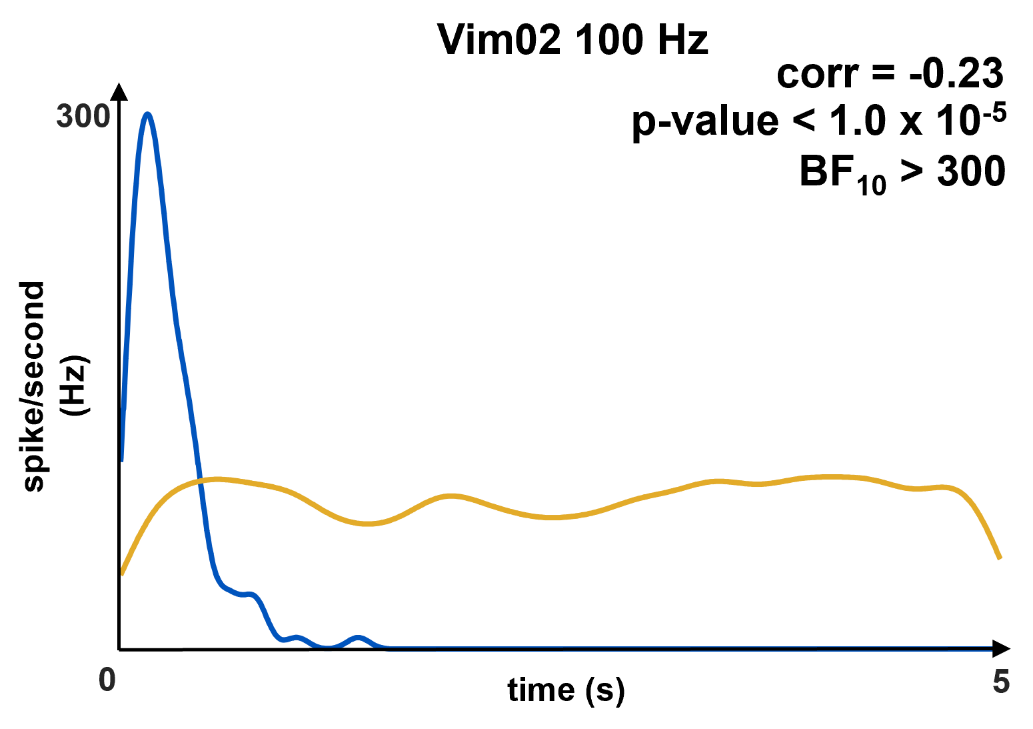

**Supplementary Figure S4 Correlation between firing rate and quasi-quasi-evoked inhibition rate:** The rate of presence of quasi-quasi-evoked inhibition is computed using an optimized width of a Gaussian kernel. This is plotted in yellow over the entire time interval. Firing rate is also computed using a width consistent with the Hideaki optimized Gaussian kernel method. This is plotted in blue over the entire time interval. Correlation, bayes factor and p-values are computed between the two curves and displayed in the top right of each plot.

**Supplementary table**

| **Neuron number** | **Available stimulation recording frequencies (Hz)** |
| --- | --- |
| Vim01 | 2, 3, 5, 10, 20, 50, 100 |
| **Vim02** | **1, 2, 3, 5, 10, 20, 50, 100** |
| Vim03 | 1, 2, 3, 5, 10, 20, 50, 100 |
| Vim04 | 1, 2, 3, 5, 10, 20, 50, 100 |
| Vim05 | 1, 2, 3, 5, 20, 50, 100 |
| Vim06 | 1, 2, 3, 5, 10, 20, 50, 100 |
| Vim07 | 1, 2, 3, 5, 10, 20, 50, 100 |
| Vim08 | 1, 2, 3, 5, 10, 20, 50, 100 |
| Vim09 | 1, 2, 3, 5, 10, 20, 50, 100 |
| **Vim10** | **100, 200** |
| **Vim11** | **100, 200** |
| Vim12 | 100, 200 |
| Vim13 | 100, 200 |
| Vim14 | 100, 200 |
| Vim15 | 200 |
| Vim16 | 100, 200 |
| Vim17 | 100, 200 |
| Vim18 | 100, 200 |
| **Vim19** | **100, 200** |

**Supplementary Table S1 Summary of recorded frequencies:** We show the neuron identifiers that recordings were taken from, each being an individual patient. We also show the stimulation frequencies that we analyzed the recordings of for each of the corresponding neuron identifiers. Rows bolded represent neurons with quasi-quasi-evoked inhibition.
